# Epidemiological transcriptomic data supports BCG protection in viral diseases including COVID-19

**DOI:** 10.1101/2020.11.10.374777

**Authors:** Abhay Sharma

## Abstract

Epidemiological and clinical evidence suggests that Bacille Calmette-Guérin (BCG) vaccine induced trained immunity protects against non-specific infections. Multiple clinical trials are currently underway to assess effectiveness of the vaccine in the coronavirus disease 2019 (COVID-19). However, the durability and mechanism of BCG trained immunity remain unclear. Here, an integrative analysis of available epidemiological transcriptomic data related to BCG vaccination and respiratory tract viral infections, and transcriptomic alterations reported in COVID-19 is presented toward addressing this gap. Results suggest that the vaccine induces very long-lasting transcriptomic changes that, unsurprisingly, mimic viral infections by upregulated antiviral defense response, and, counterintuitively. oppose viral infections by downregulated myeloid cell activation. These durability and mechanistic insights have immediate implications in fight against the COVID-19 pandemic.

## Introduction

Bacille Calmette-Guérin (BCG) vaccine, besides conferring adaptive immunity against tuberculosis, is considered to induce trained immunity, a general long-term stimulation of innate immune response that provides nonspecific protection against pathogens, especially respiratory tract viral infections (*1–3*). Insights from human vaccination and experimental infection models, and animal models suggest that transcriptional, epigenetic, and functional reprogramming of hematopoietic stem cells toward myelopoiesis, and epigenetic modifications of peripheral myeloid cells drive BCG induced trained immunity (*4–7*). Currently, the prospect that the vaccine may limit the impact of the coronavirus disease 2019 (COVID-19) pandemic is a subject of immense interest (*8–12*). Epidemiological studies have shown that countries with childhood BCG vaccination policy, compared to nations without the policy, faced lower prevalence and mortality of COVID-19 (*13, 14*). However, at the same time,, no association between childhood vaccination and protection from COVID-19 in adulthood has also been reported (*15*), fueling the speculation that BCG induced trained immunity might be short-lived (*16*). Meanwhile, a recent retrospective study of a cohort of adults who received or did not receive the vaccine in the preceding 5 years has suggested that BCG vaccination is possibly associated with a lower incidence of illness during the pandemic (*16*). Also, while interventional clinical trials are underway to test whether BCG may have an immediate beneficial effect in COVID-19, a randomized clinical trial has shown that BCG vaccination in the elderly lengthens the time to first infection and reduces the incidence of new infection, especially of respiratory tract viruses (*3*). Interestingly, both the above studies (*3, 16*) surprisingly found no evidence of excessive systemic inflammation in vaccinated subjects, alleviating the concern that BCG trained immunity, due to elevated innate immune response, may add to hyperinflammation in severe disease and worsen the condition (*3*). The negative evidence of inflammation has been indirectly explained by arguing that BCG vaccination may lower systemic inflammation through enhanced antiviral defense response leading to decreased viral loads (*3, 16*). Taken together, an improved understanding of both the durability and the mechanisms of BCG induced trained immunity is urgently required, particularly in view of the concern that indiscriminate use of the vaccine may jeopardize supply required for vaccination of children to protect them against tuberculosis, and the possibility that BCG may produce deleterious effects in severe COVID-19 patients (*10*). Given the potential for knowledge discovery from themed collection and integration of available transcriptomic datasets (*17*), an integrative analysis of human epidemiological transcriptomic data related to BCG vaccination and viral respiratory infections, along with COVID-19 transcriptomic data, is presented here toward understanding the durability and mechanism of trained immunity.

## Results

Clustering differentially expressed genes (DEG) in whole blood (WB) of infants 2, 6, 12, 18 and 26 weeks following BCG vaccination (*18*), compared to the baseline, show directionally consistent expression changes across time points (**Figure 1A**). Similarly, clustering DEG in BCG stimulated peripheral blood mononuclear cells (PBMC), compared to unstimulated PBMC, of BCG naïve adults and BCG vaccinated adults - median time since vaccination, 10 years - before and after 2, 7 and 14 days of an intradermal BCG challenge (*19*) shows directionally consistent changes across time points in general, with vaccinated subjects exhibiting higher level of transcriptomic alterations than naïve subjects (**Figure 1B**). Notably, clustering of common genes between the above two clusters (**Figure 1A, B**) shows that genes that are up- or down-regulated in WB weeks and months after vaccination are similarly regulated in PBMC after BCG stimulation, more so in vaccinated individuals (**Figure 1C**). These results clearly suggest that, consistent with the concept of trained immunity, BCG vaccination leads to persistent changes in peripheral blood cell transcriptome. Next, clustering of DEG in WB of patients with varying severity and time points, and in- and out-patient status of illness from infection with respiratory viruses (*20–23*), compared to healthy or appropriate patient controls, shows transcriptomic changes that are both common across viruses as well as virus specific, as expected, and differ in magnitude in consonance with the variables (**Figure 1D**). Remarkably, clustering of DEG that are common between the short-term BCG group (**Figure 1A**) and virus group (**Figure 1D**) shows strikingly similar and contrasting pattern of altered gene expression between the BCG and the virus group as a whole; (i) a major set of genes that are upregulated in both the groups, (ii) another major set of genes that are downregulated in the BCG and upregulated in the virus group, (iii) a minor set of genes that are downregulated in both the groups, and (iv) another minor set of genes that are upregulated in the BCG and downregulated in the virus group (**Figure 1E**). Merging long-term BCG group (**Figure 1B**) further tightened this pattern (**Figure 1F**). The gene regulation observed in the virus group (**Figure 1F**) is validated in general by an independent cluster of DEG in WB or PBMC of adults, infants, and children with infection of various respiratory viruses (*24–26*), compared to healthy controls (**Figure 1G**). Further independent validation is obtained by clustering DEG in WB of adult patients in a cohort of acute respiratory illness (ARI) from infection with various viruses (*27*), compared to the baseline (**Figure 1H**); with day 0, representing up to 48 hours of ARI onset, resembling gene regulation observed in the virus group (**Figure 1F**), and later time points showing absence of differential expression of genes that were upregulated by both the BCG and the virus group (**Figure 1F**), and reversal from up- to down-regulation for genes that were upregulated in the virus group and downregulated in the BCG group. Another independent validation is observed with adult patient-specific WB pattern of gene expression in COVID-19 (*28*); z-score transformed and normalized gene counts in severe patient, especially at time points of higher supplemental oxygen, resembling the virus group (**Figure 1F**), and in the same patient during recovery or in mild patients showing a trend toward opposite regulation (**Figure 1I**). Gene ontology analysis of all the genes in the final BCG-virus cluster (**Figure 1F**) shows enrichment of various biological process terms (**Figure 1J**); mainly, the genes that are upregulated in both the groups are associated with terms related to type I interferon signaling, cytokine-mediated signaling, and defense response to virus, whereas genes that are upregulated in the virus group and downregulated in the BCG group are associated with myeloid leukocyte activation, myeloid leukocyte mediated immunity, and other related terms. Next, genes in the final BCG-virus cluster (**Figure 1F**) were examined for blood cell type distribution, with the results demonstrating a clear myeloid bias for genes that are downregulated in the BCG and upregulated in the virus group, and no gross bias for genes that are upregulated in both the groups (**Figure 1K**). Finally, to validate this finding in disease setting, the BCG-virus cluster genes (**Figure 1F**) were intersected with DEG identified in single-cell RNA-seq analysis of PBMC from severe and severe-moderate patients of COVID-19 (*29, 30*), compared to healthy controls; as expected from the above cell type distribution analysis (**Figure 1F**), a clear myeloid bias was found for the genes that are downregulated in the BCG and upregulated in the virus group, whereas genes that are upregulated in both the groups did not starkly differ in myeloid-lymphoid distribution (**Figure 1L**). Also, unlike severe patients, severe-moderate cases did not show a clear upregulation of genes that are downregulated in the BCG and upregulated in the virus group (**Figure 1F**), consistent with observed differences (**Figure 1D-F, H, I**) among patients with varying disease status including severity and time points.

**Figure 1.**
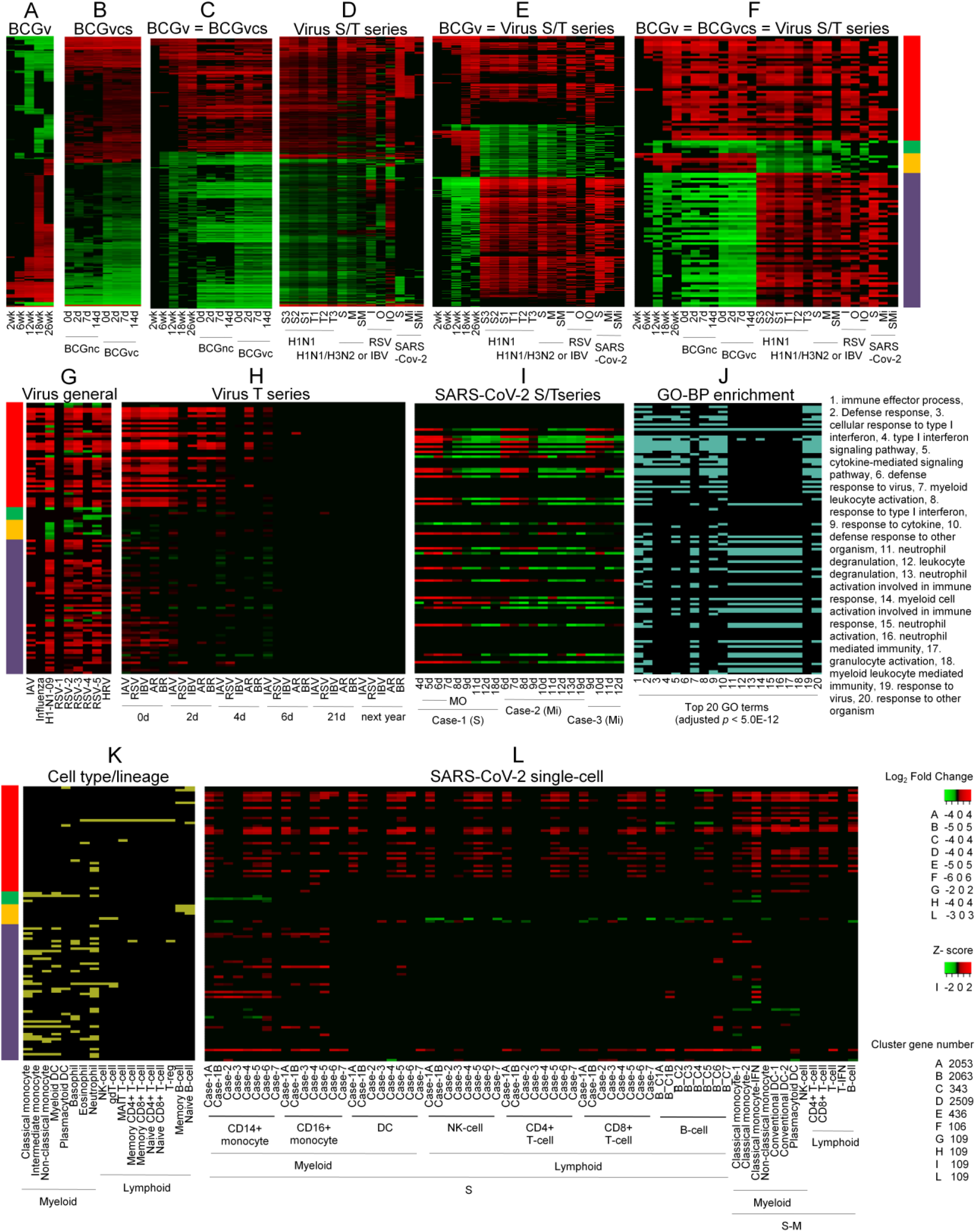
Integrative transcriptomics of BCG vaccination and viral respiratory diseases. **(A-I)** Clustering of DEG identified under indicated conditions. Abbreviation: BCGv, BCG vaccination; BCGvcs, BCG vaccination, challenge, and stimulation; BCGnc, BCG naïve and challenged; BCGvc, BCG vaccination and challenged; S/T series, both severity and time series; S3-S1, severity in decreasing order; T1-T3, sequentially from time of patient enrolment to clinical resolution; S, severe; M, moderate; SM, severe compared to moderate; I, inpatient; O, outpatient; IO, inpatient compared to outpatient; Mi, mild; SMi, severe compared to mild; IBV, influenza B virus; RSV, respiratory syncytial virus; IAV, influenza A virus, RSV-1 to −5, five different RSV datasets; HRV, human rhinovirus; T series, time series; AR, both IAV and HRV; BR, both ABV and HRV; MO, maximum supplemental oxygen. Clusters BCGvcs (B) and Virus S/T series (D) are composed of most frequent DEG among samples. = indicates common genes in the given clusters. Multicolored bar represent DEG directionality: red (42 genes), upregulation in all BCG and all virus samples; green (5 genes), downregulation in all BCG and all virus samples; orange (8 genes), upregulation in all BCG samples and downregulation in all virus samples; purple (54 genes), downregulation in all BCG samples and upregulation in virus samples. Sample specific details including original reference and data source, DEG identification method, and cluster values are indicated in **Supplementary Table 1**. (**J**) Gene ontology biological process (GO-BP) enrichment of genes in the final BCG-virus cluster (F, BCGv = BCGvcs = Virus S/T series). Full results are presented in **Supplementary Table 2**. (**K**) Distribution of genes in BCGv = BCGvcs = Virus S/T series cluster (F) among cell type elevated genes. Distribution in tabular form is provided in **Supplementary Table 3**. (**L**) Clustering of DEG identified in single cell RNA-seq analysis of severe (S) and severe-moderate (S-M) COVID-19 patients. Matching cell types in S and S-M are shown. Sample specific details including original reference and data source, DEG identification method, and cluster values are indicated in **Supplementary Table 4**.

## Discussion

The present analysis suggests that BCG vaccination induces long-term changes in blood immune cell transcriptome that partly mimic and partly oppose transcriptomic changes induced by viral respiratory illnesses including COVID-19. The mimicking part relates to upregulation of defense response to virus, and the opposing part to downregulation of myeloid cell activation in vaccinated subjects, with upregulation of the same observed in patients. Given these, individuals vaccinated with BCG in the near or distant past would be expected to have a higher basal level of antiviral defense and a lower basal level of myeloid cell activation, rendering them less susceptible to viral infections on account of the former, and to hyperinflammation, that characterizes severe disease (*16*), on account of the latter. This prediction is consistent with the epidemiological studies showing lower prevalence and mortality of COVID-19 in countries with childhood BCG vaccination policy (*13*). Regarding mechanism, the present finding is in line with the existing concept (*2*) that trained immunity involves reprogramming of myeloid cells, and enhanced capacity for cytokine production and antimicrobial function. However, instead of myeloid cell activation genes showing upregulation in vaccinated subjects, as might have been possibly expected on account of enhanced innate immune response that is considered to characterize trained immunity, these genes show persistent downregulation postvaccination. Notably, a retrospective cohort study (*16*) and a randomized clinical trial (*3*) have recently reported that BCG vaccination is not associated with systemic inflammation, alleviating the concern that the vaccine may produce deleterious effects in severe COVID-19 by adding to hyperinflammation. That BCG is not associated with inflammation (*3, 16*) has been indirectly explained by arguing that BCG induced enhancement in antiviral defense would cause decreased viral loads which in turn would lead to lower systemic inflammation (*16*). In contrast, the analysis presented here would suggest that a suppressed level of myeloid cell activation may directly limit innate immunity response from causing inflammation. In conclusion, the present evidence suggesting high durability of BCG induced trained immunity provides a rationale against possible indiscriminate use (*10*) of BCG vaccination as a measure to reduce the impact of COVID-19 pandemic. Also, it offers a mechanistic reasoning to ease the concern (*10*) that BCG induced trained immunity may escalate systemic inflammation and worsen the condition in severe COVID-19 patients.

## Materials and methods

Relevant publications and datasets were manually identified by extensively searching NCBI’s PubMed and Gene Expression Omnibus (GEO), with a preference for large and comparative studies. Original author-identified gene sets were used, if available in full. Otherwise, differentially expressed genes (DEG) with adjusted *p* value significance were identified from GEO datasets by using GEO2R (*31*) for microarray or GREIN (*32*) for RNA-seq studies. DEG were clustered and heatmaps produced using Heatmapper (*33*), with, wherever applicable, scale type none, and average linkage and Euclidean distance. Gene ontology biological process enrichment was determined using ToppGene Suite (*34*). The list of cell type elevated genes in blood atlas (*35*) was used for cell type distribution analysis.

## Data availability

All data is available in the manuscript or the supplementary materials.

## References

1. M. G. Netea, L. A. B. Joosten, E. Latz, K. H. G. Mills, G. Natoli, H. G. Stunnenberg, L. A. J. O’Neill, R. J. Xavier, Trained immunity: A program of innate immune memory in health and disease. Science 352, aaf1098 (2016).

2. M. G. Netea, J. Domínguez-Andrés, L. B. Barreiro, T. Chavakis, M. Divangahi, E. Fuchs, L. A. B. Joosten, J. W. M. van der Meer, M. M. Mhlanga, W. J. M. Mulder, N. P. Riksen, A. Schlitzer, J. L. Schultze, C. S. Benn, J. C. Sun, R. J. Xavier, E. Latz, Defining trained immunity and its role in health and disease. Nat. Rev. Immunol. 20, 375–388 (2020).

3. E. J. Giamarellos-Bourboulis, M. Tsilika, S. Moorlag, N. Antonakos, A. Kotsaki, J. Domínguez-Andrés, E. Kyriazopoulou, T. Gkavogianni, M. E. Adami, G. Damoraki, P. Koufargyris, A. Karageorgos, A. Bolanou, H. Koenen, R. van Crevel, D. I. Droggiti, G. Renieris, A. Papadopoulos, M. G. Netea, Activate: Randomized Clinical Trial of BCG Vaccination against Infection in the Elderly. Cell 183, 315–323 (2020).

4. E. Kaufmann, J. Sanz, J. L. Dunn, N. Khan, L. E. Mendonça, A. Pacis, F. Tzelepis, E. Pernet, A. Dumaine, J. C. Grenier, F. Mailhot-Léonard, E. Ahmed, J. Belle, R. Besla, B. Mazer, I. L. King, A. Nijnik, C. S. Robbins, L. B. Barreiro, M. Divangahi, BCG educates hematopoietic stem cells to generate protective innate immunity against tuberculosis. Cell 172, 176–190 (2018).

5. B. Cirovic, L. C. J. de Bree, L. Groh, B. A. Blok, J. Chan, W. J. F. M. van der Velden, M. E. J. Bremmers, R. van Crevel, K. Händler, S. Picelli, J. Schulte-Schrepping, K. Klee, M. Oosting, V. A. C. M. Koeken, J. van Ingen, Y. Li, C. S. Benn, J. L. Schultze, L. A. B. Joosten, N. Curtis, M. G. Netea, A. Schlitzer, BCG vaccination in humans elicits trained immunity via the hematopoietic progenitor compartment. Cell Host Microbe 28, 322–334 (2020).

6. N. Khan, J. Downey, J. Sanz, E. Kaufmann, B. Blankenhaus, A. Pacis, E. Pernet, E. Ahmed, S. Cardoso, A. Nijnik, B. Mazer, C. Sassetti, M. A. Behr, M. P. Soares, L. B. Barreiro, M. Divangahi, M. tuberculosis reprograms hematopoietic stem cells to limit myelopoiesis and impair trained immunity. Cell 183, 752–770 (2020).

7. R. J. W. Arts, S. J. C. F. M. Moorlag, B. Novakovic, Y. Li, S. Y. Wang, M. Oosting, V. Kumar, R. J. Xavier, C. Wijmenga, L. A. B. Joosten, C. B. E. M. Reusken, C. S. Benn, P. Aaby, M. P. Koopmans, H. G. Stunnenberg, R. van Crevel, M. G. Netea, BCG Vaccination Protects against Experimental Viral Infection in Humans through the Induction of Cytokines Associated with Trained Immunity. Cell Host Microbe 23, 89–100 (2018).

8. L. A. J. O’Neill, M. G. Netea, BCG-induced trained immunity: can it offer protection against COVID-19? Nat. Rev. Immunol. 20, 335–337 (2020).

9. A. Mantovani, M. G. Netea, Trained Innate Immunity, Epigenetics, and Covid-19. N. Engl. J. Med. 383, 1078–1080 (2020).

10. N. Curtis, A. Sparrow, T. A. Ghebreyesus, M. G. Netea, Considering BCG vaccination to reduce the impact of COVID-19. Lancet 395, 1545–1546 (2020).

11. M. G. Netea, E. J. Giamarellos-Bourboulis, J. Domínguez-Andrés, N. Curtis, R. van Crevel, F. L. van de Veerdonk, M. Bonten, Trained Immunity: a Tool for Reducing Susceptibility to and the Severity of SARS-CoV-2 Infection. Cell 181, 969–977 (2020).

12. G, Redelman-Sidi, Could BCG be used to protect against COVID-19? Nat. Rev. Urol. 17, 316–317 (2020).

13. M. K. Berg, Q. Yu, C. E. Salvador, I. Melani, S. Kitayama, Mandated Bacillus Calmette-Guérin (BCG) vaccination predicts flattened curves for the spread of COVID-19. Sci. Adv. 6, eabc1463 (2020).

14. L. E. Escobar, A. Molina-Cruz, C. Barillas-Mury, BCG vaccine protection from severe coronavirus disease 2019 (COVID-19). Proc. Natl. Acad. Sci. U S A 117, 17720–17726 (2020).

15. U. Hamiel, E. Kozar, I. Youngster, SARS-CoV-2 Rates in BCG-Vaccinated and Unvaccinated Young Adults. JAMA 323, 2340–2341 (2020).

16. S. J. C. F. M. Moorlag, R. C. van Deuren, C. H. van Werkhoven, M. Jaeger, P. Debisarun, E. Taks, V. P. Mourits, V. A. C. M. Koeken, L. C. J. de Bree, T. Doesschate, M. C. Cleophas, S. Smeekens, M. Oosting, F. L. van de Veerdonk, L. A. B. Joosten, J. T. Oever, J. W. M. van der Meer, N. Curtis, P. Aaby, C. Stabell-Benn, E. J. Giamarellos-Bourboulis, M. Bonten, R. van Crevel, M. G. Netea, Safety and COVID-19 Symptoms in Individuals Recently Vaccinated with BCG: a Retrospective Cohort Study. Cell Rep. Med. 1, 100073 (2020).

17. Z. Wang, A. Lachmann, A. Ma’ayan, Mining data and metadata from the gene expression omnibus. Biophys. Rev. 11, 103–110 (2019).

18. A. G. Loxton, J. K. Knaul, L. Grode, A. Gutschmidt, C. Meller, B. Eisele, H. Johnstone, G. van der Spuy, J. Maertzdorf, S. H. E. Kaufmann, A. C. Hesseling, G. Walzl, M. F. Cotton, Safety and Immunogenicity of the Recombinant Mycobacterium bovis BCG Vaccine VPM1002 in HIV-Unexposed Newborn Infants in South Africa. Clin. Vaccine Immunol. 24, e00439–16 (2017).

19. M. Matsumiya, I. Satti, A. Chomka, S. A. Harris, L. Stockdale, J. Meyer, H. A. Fletcher, H. McShane, Gene expression and cytokine profile correlate with mycobacterial growth in a human BCG challenge model. J. Infect. Dis. 211, 1499–1509 (2015).

20. J. Dunning, S. Blankley, L. T. Hoang, M. Cox, C. M. Graham, P. L. James, C. I. Bloom, D. Chaussabel, J. Banchereau, S. J. Brett, M. F. Moffatt, A. O’Garra, P. J. M. Openshaw; MOSAIC Investigators, Progression of whole-blood transcriptional signatures from interferon-induced to neutrophil-associated patterns in severe influenza. Nat. Immunol. 19, 625–635 (2018).

21. B. M. Tang, M. Shojaei, S. Teoh, A. Meyers, J. Ho, T. B. Ball, Y. Keynan, A. Pisipati, A. Kumar, D. P. Eisen, K. Lai, M. Gillett, R. Santram, R. Geffers, J. Schreiber, K. Mozhui, S. Huang, G. P. Parnell, M. Nalos, M. Holubova, T. Chew, D. Booth, A. Kumar, A. McLean, K. Schughart, Neutrophils-related host factors associated with severe disease and fatality in patients with influenza infection. Nat. Commun. 10, 3422 (2019).

22. W. A. de Steenhuijsen Piters, S. Heinonen, R. Hasrat, E. Bunsow, B. Smith, M. C. Suarez-Arrabal, D. Chaussabel, D. M. Cohen, E. A. Sanders, O. Ramilo, D. Bogaert, A. Mejias, Nasopharyngeal microbiota, host transcriptome, and disease severity in children with respiratory syncytial virus infection. Am. J. Respir. Crit. Care Med. 194, 1104–1115 (2016).

23. A. C. Aschenbrenner, M. Mouktaroudi, B. Kraemer, N. Antonakos, M. Oestreich, K. Gkizeli, M. Nuesch-Germano, M. Saridaki, L. Bonaguro, N. Reusch, K. Bassler, S. Doulou, R. Knoll, T. Pecht, T. S. Kapellos, N. Rovina, C. Kroeger, M. Herbert, L. Holsten, A. Horne, I. D. Gemuend, S. Agrawal, K. Dahm, M. van Uelft, A. Drews, L. Lenkeit, N. Bruse, J. Gerretsen, J. Gierlich, M. Becker, K. Haendler, M. Kraut, H. Theis, S. Mengiste, E. D. Domenico, J. Schulte-Schrepping, L. Seep, J. Raabe, C. Hoffmeister, M. ToVinh, V. Keitel, G. J. Rieke, V. Talevi, A. N. Aziz, P. Pickkers, F. van de Veerdonk, M. G. Netea, J. L. Schultze, M. Kox, M. M. B. Breteler, J. Nattermann, A. Koutsoukou, E. J. Giamarellos-Bourboulis, T. Ulas, Disease severity-specific neutrophil signatures in blood transcriptomes stratify COVID-19 patients. medRxiv 2020.07.07.20148395.

24. A. Mejias, B. Dimo, N. M. Suarez, C. Garcia, M. C. Suarez-Arrabal, T. Jartti, D. Blankenship, A. Jordan-Villegas, M. I. Ardura, Z. Xu, J. Banchereau, D. Chaussabel, O. Ramilo, Whole blood gene expression profiles to assess pathogenesis and disease severity in infants with respiratory syncytial virus infection. PLoS Med. 10, e1001549 (2013).

25. I. Ioannidis, B. McNally, M. Willette, M. E. Peeples, D. Chaussabel, J. E. Durbin, O. Ramilo, A. Mejias, E. Flaño, Plasticity and virus specificity of the airway epithelial cell immune response during respiratory virus infection. J. Virol. 86, 5422–5436 (2012).

26. J. A. Herberg, M. Kaforou, S. Gormley, E. R. Sumner, S. Patel, K. D. Jones, S. Paulus, C. Fink, F. Martinon-Torres, G. Montana, V. J. Wright, M. Levin, Transcriptomic profiling in childhood H1N1/09 influenza reveals reduced expression of protein synthesis genes. J. Infect. Dis. 208, 1664–1668 (2013).

27. Y. Zhai, L. M. Franco, R. L. Atmar, J. M. Quarles, N. Arden, K. L. Bucasas, J. M. Wells, D. Niño, X. Wang, G. E. Zapata, C. A. Shaw, J. W. Belmont, R. B. Couch, Host Transcriptional Response to Influenza and Other Acute Respiratory Viral Infections--A Prospective Cohort Study. PLoS Pathog. 11, e1004869 (2015).

28. E. Z. Ong, Y. F. Z. Chan, W. L. Leong, N. M. Y. Lee, S. Kalimuddin, S. M. Haja Mohideen, K. S. Chan, A. T. Taa, A. Bertoletti, E. E. Ooi, J. G. H. Low, A Dynamic Immune Response Shapes COVID-19 Progression. Cell Host Microbe 27, 879–882 (2020).

29. A. J. Wilk, A. Rustagi, N. Q. Zhao, J. Roque, G. J. Martínez-Colón, J. McKechnie, G. T. Ivison, T. Ranganath, R. Vergara, T. Hollis, L. J. Simpson, P. Grant, A. Subramanian, A. J. Rogers, C. A. Blish, A single-cell atlas of the peripheral immune response in patients with severe COVID-19. Nat. Med. 26, 1070–1076 (2020).

30. P. S. Arunachalam, F. Wimmers, C. K. P. Mok, R. A. P. M. Perera, M. Scott, T. Hagan, N. Sigal, Y. Feng, L. Bristow, O. Tak-Yin Tsang, D. Wagh, J. Coller, K. L. Pellegrini, D. Kazmin, G. Alaaeddine, W. S. Leung, J. M. C. Chan, T. S. H. Chik, C. Y. C. Choi, C. Huerta, M. Paine McCullough, H. Lv, E. Anderson, S. Edupuganti, A. A. Upadhyay, S. E. Bosinger, H. T. Maecker, P. Khatri, N. Rouphael, M. Peiris, B. Pulendran, Systems biological assessment of immunity to mild versus severe COVID-19 infection in humans. Science 369, 1210–1220 (2020).

31. T. Barrett, S. E. Wilhite, P. Ledoux, C. Evangelista, I. F. Kim, M. Tomashevsky, K. A. Marshall, K. H. Phillippy, P. M. Sherman, M. Holko, A. Yefanov, H. Lee, N. Zhang, C. L. Robertson, N. Serova, S. Davis, A. Soboleva, NCBI GEO: archive for functional genomics data sets--update. Nucleic Acids Res. 41(Database issue), D991–D995 (2013).

32. N. A. Mahi, M. F. Najafabadi, M. Pilarczyk, M. Kouril, M. Medvedovic, GREIN: An Interactive Web Platform for Re-analyzing GEO RNA-seq Data. Sci. Rep. 9, 7580 (2019).

33. S. Babicki, D. Arndt, A. Marcu, Y. Liang, J. R. Grant, M. Maciejewski, D. S. Wishart, Heatmapper: web-enabled heat mapping for all. Nucleic Acids Res. 44(W1), W147–W153 (2016).

34. J. Chen, E. E. Bardes, B. J. Aronow, A. G. Jegga, ToppGene Suite for gene list enrichment analysis and candidate gene prioritization. Nucleic Acids Res. 37(Web Server issue), W305–W311 (2009).

35. M. Uhlen, M. J. Karlsson, W. Zhong, A. Tebani, C. Pou, J. Mikes, T. Lakshmikanth, B. Forsström, F. Edfors, J. Odeberg, A. Mardinoglu, C. Zhang, K. von Feilitzen, J. Mulder, E. Sjöstedt, A. Hober, P. Oksvold, M. Zwahlen, F. Ponten, C. Lindskog, A. Sivertsson, L. Fagerberg, P. Brodin, A genome-wide transcriptomic analysis of protein-coding genes in human blood cells. Science 366, eaax9198 (2019).

